# An Age-Specific Atlas for Delineation of White Matter Pathways in Children Aged 6-8 Years

**DOI:** 10.1101/2020.06.21.157222

**Authors:** Arthur P.C. Spencer, Hollie Byrne, Richard Lee-Kelland, Sally Jary, Marianne Thoresen, Frances M. Cowan, Ela Chakkarapani, Jonathan C.W. Brooks

**Author notes:** Corresponding author; Clinical Research and Imaging Centre, University of Bristol, 60 St Michael’s Hill, Bristol, BS2 8DX.

## Abstract

Diffusion MRI allows non-invasive assessment of white matter maturation in typical development and of white matter damage due to brain injury or pathology. Probabilistic white matter atlases provide delineation of white matter tracts, allowing diffusion metrics to be measured in specific white matter pathways. However, given the known age-dependency of developmental change in white matter it may not be optimal to use an adult template when assessing data acquired from children. This study develops an age-specific probabilistic white matter atlas for delineation of 12 major white matter tracts in children aged 6-8 years. By comparing to subject-specific tract tracing in two validation cohorts, we demonstrate that this age-specific atlas gives better overall performance than simply registering to the Johns Hopkins University adult white matter template. Specifically, when normalising diffusion data acquired from children to an adult template, estimates of fractional anisotropy (FA) values for corticospinal tract, uncinate fasciculus, forceps minor, cingulate gyrus part of the cingulum and anterior thalamic radiation were all less accurate than those obtained when using an age-specific atlas, potentially leading to false negatives when performing group comparisons. We then applied the newly developed atlas to compare FA between children treated with therapeutic hypothermia for neonatal encephalopathy and age-matched controls, which revealed significant reductions in the fornix, the left superior longitudinal fasciculus, and both the hippocampal and cingulum parts of the left cingulate gyrus. To our knowledge, this is the first publicly available probabilistic atlas of white matter tracts for this age group.

## 1 Introduction

Tract-level analysis of diffusion weighted imaging (DWI) data is used extensively to investigate white matter microstructure in both typical (Asato et al., 2010; Hüppi and Dubois, 2006; Lebel et al., 2008) and atypical brain development (for a review, see (Dennis and Thompson, 2013)). In children and adolescents, atypical brain development may lead to physical and intellectual disabilities including e.g. cerebral palsy (CP) (Arrigoni et al., 2016), autistic spectrum behaviours (Ameis and Catani, 2015; Dimond et al., 2019) and attention deficit hyperactivity disorder (Konrad and Eickhoff, 2010). Diffusion metrics such as fractional anisotropy (FA), mean diffusivity (MD), radial diffusivity (RD) and axial diffusivity (AD) (Assaf and Pasternak, 2008) are sensitive to changes in the underlying white matter structure. These metrics are widely investigated in studies of brain development (Dennis and Thompson, 2013; Lebel et al., 2008), as well as having clinical relevance in patient cohorts (Assaf et al., 2019; Assaf and Pasternak, 2008; Horsfield and Jones, 2002).

To measure tract-level diffusion metrics, white matter tracts can be delineated by registering to a standard template with a probabilistic atlas of tract locations. Using a white matter atlas eliminates the need for computationally intensive methods of delineating tracts by segmenting streamlines generated by tractography (Lawes et al., 2008; Sydnor et al., 2018; Wakana et al., 2007; Wassermann et al., 2010; Zhang et al., 2018). This is beneficial in clinical settings or when studying large datasets. Additionally, data which have been acquired with shorter, more simplistic diffusion tensor acquisitions may not facilitate accurate tractography. Such acquisitions may be favoured in an effort to minimise scan times (and therefore minimise risk of movement during the scan) when studying children, including those with disabilities who would benefit from investigating white matter development (Phan et al., 2018).

The widely used Johns Hopkins University (JHU) white matter tract atlas (Hua et al., 2008) is constructed from adult data. Numerous developmental studies demonstrate white matter alterations continuing into adolescence (Cascio et al., 2007; Hagmann et al., 2010; Lebel et al., 2008; Simmonds et al., 2014), and white matter development varies widely across the brain (Lebel et al., 2019), therefore an atlas constructed from adult scans is by design and definition not representative of children. There are several publicly available age-specific structural templates (Altaye et al., 2008; Fonov et al., 2011; Richards et al., 2016; Sanchez et al., 2012), however none of these provide diffusion data.

Using robust tract reconstruction protocols (Hua et al., 2008; Wakana et al., 2007) this study develops an age-specific probabilistic white matter atlas for 12 major tracts in children aged 6-8 years. To assess whether this atlas accurately delineates tracts, we measured both volumetric overlap and the diffusion metrics sampled by the tract mask in comparison with subject-specific tractography-based tract delineation. We then assess the utility of this age-specific tract atlas by comparing it to results obtained using an adult atlas (JHU). The atlas is then further validated against an open data source (i.e. different scanner and acquisition protocol), and against a different tractography algorithm.

As a demonstration, we then investigate tract-level differences in children treated with therapeutic hypothermia (TH) for neonatal encephalopathy (NE) at birth, compared with healthy controls, and compare results obtained using the age-specific atlas to those from the JHU atlas. The children who had TH, do not have CP and are in mainstream education still exhibit significantly reduced performance on cognitive tests (Jary et al., 2019; Lee-Kelland et al., 2020) and have slower reaction times and reduced visuo-spatial processing abilities (Tonks et al., 2019), compared to the typically developing controls.

## 2 Material and Methods

### 2.1 Participants

Ethics approval was obtained from the North Bristol Research Ethics Committee and the Health Research Authority (REC ID: 15/SW/0148). Informed and written consent was obtained from the parents of participants before collecting data. The cohort was made up of 36 healthy children aged 6-8 years with no evidence of neurological disease, originally recruited as controls for a study of the long-term effects of TH (“CoolMRI”) on behavioural and imaging outcomes. The 36 controls were split randomly into 28 atlas and 8 validation subjects such that the group were matched for age, sex, socio-economic status (SES) and full-scale intelligence quotient (FSIQ). For the demonstrative case study, data from 33 children treated with TH following NE at birth were compared to the control data.

### 2.2 Image Acquisition

DWI data were acquired with a Siemens 3 tesla Magnetom Skyra MRI scanner at the Clinical Research and Imaging Centre (CRiCBristol), Bristol, UK. Subjects were placed supine within the 32-channel receive only head-coil by an experienced radiographer, and head movement minimised by means of memory-foam padding. Children wore earplugs and were able to watch a film. A multiband echo-planar imaging (EPI) sequence was used with the following parameters: TE = 70 ms; TR = 3150 ms; field of view 192 × 192 mm; 60 slices; 2.0 mm isotropic voxels; phase encoding in the anterior-posterior direction, in-plane acceleration factor = 2 (GRAPPA (Griswold et al., 2002)), through-plane multi-band factor = 2 (Moeller et al., 2010; Setsompop et al., 2012b, 2012a).

For the purpose of data averaging and eddy-current distortion correction, two sets of diffusion weighted images were acquired with b = 1,000 s mm^-2^ in 60 diffusion directions, equally distributed according to an electrostatic repulsion model, as well as 8 interspersed b = 0 images, with one data set acquired with positive phase encoding steps, then repeated with negative steps (so-called, “blip-up”, “blip-down”), giving a total of 136 images.

### 2.3 Quality Control

The quality of the diffusion data was assessed using the EddyQC tool (Bastiani et al., 2019) from FSL (Smith et al., 2004). This provides several measures of the amount of movement and eddy current induced distortion present in the data. For each participant, metrics were normalised, then the root-mean-square was calculated, giving a score which increases monotonically with the amount of movement and eddy current distortion. Scans were rejected if their score was more than one standard deviation above the mean of all participants.

### 2.4 Image Processing & Atlas Construction

DWI data were corrected for eddy current induced distortions and subject movements using EDDY (Andersson and Sotiropoulos, 2016) and TOPUP (Andersson et al., 2003), part of FSL. Subsequent DWI processing and tractography steps were performed using MRtrix (Tournier et al., 2019). The response function (the DWI signal for a typical fibre population) was estimated from the data (Tournier et al., 2013). The fibre orientation distribution (FOD) was then calculated by performing constrained spherical deconvolution of the response function from the measured DWI signal (Tournier et al., 2007). Deterministic tractography was run in each subject using the “SD Stream” algorithm (Tournier et al., 2012). Streamlines were seeded randomly in the brain and generated with a step size of 0.2 mm, then terminated if the FOD amplitude dropped below 0.2 or the angle between successive steps exceeded 40 degrees. 10 million streamlines were generated, which were then filtered to 1 million using spherical-deconvolution informed filtering of tractograms (SIFT) (Smith et al., 2013) to give better reconstruction of FODs, improving anatomical accuracy.

The process of generating probability maps from the whole-brain tractograms is summarised in Figure 1. White matter tracts were segmented from whole-brain tractograms using the protocols described in Wakana et al., whereby regions of interest (ROI) are drawn to include or exclude streamlines passing through them (Wakana et al., 2007). For a given tract, any streamlines which pass through all inclusion ROIs and no exclusion ROIs belong to that tract, and all other streamlines are removed. Inclusion and exclusion ROIs were manually drawn in each subject to delineate 12 major fibre tracts: anterior thalamic radiation (ATR); cingulate gyrus part of the cingulum (CG); hippocampal part of the cingulum (CH); cortico-spinal tract (CST); forceps major (Fmajor); forceps minor (Fminor); inferior fronto-occipital fasciculus (IFOF); inferior longitudinal fasciculus (ILF); superior longitudinal fasciculus (SLF); temporal projections of the superior longitudinal fasciculus (SLFt); uncinate fasciculus (UF); and the fornix. The locations of ROIs for all tracts apart from the fornix are described in Wakana et al. as shown in Figure 2 (Hua et al., 2008; Wakana et al., 2007).

**Figure 1.**
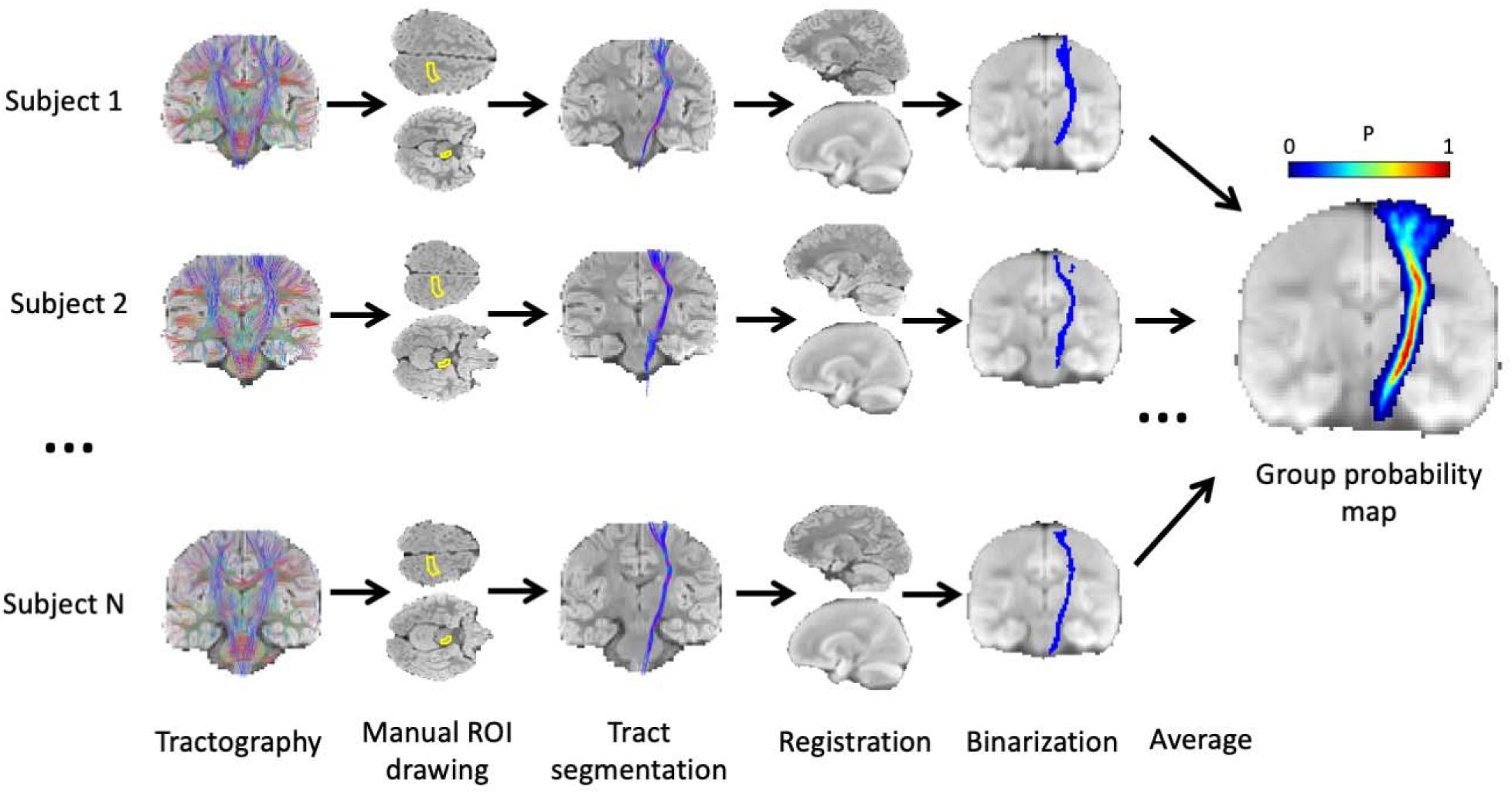
Methodology for generating probabilistic tract maps from whole-brain tractography data, shown here for the corticospinal tract (CST). ROIs were manually drawn in each subject, as defined by (Wakana et al., 2007) (in the case of the CST, inclusion ROIs were drawn in the cerebral peduncle and the primary motor cortex), and tracts were segmented by including streamlines passing through the inclusion ROIs. Streamlines were transformed to standard space (JHU template) and a binary mask was created for each subject indicating all voxels containing streamlines. The average of these masks (across N = 28 subjects) gives the probability map.

**Figure 2.**
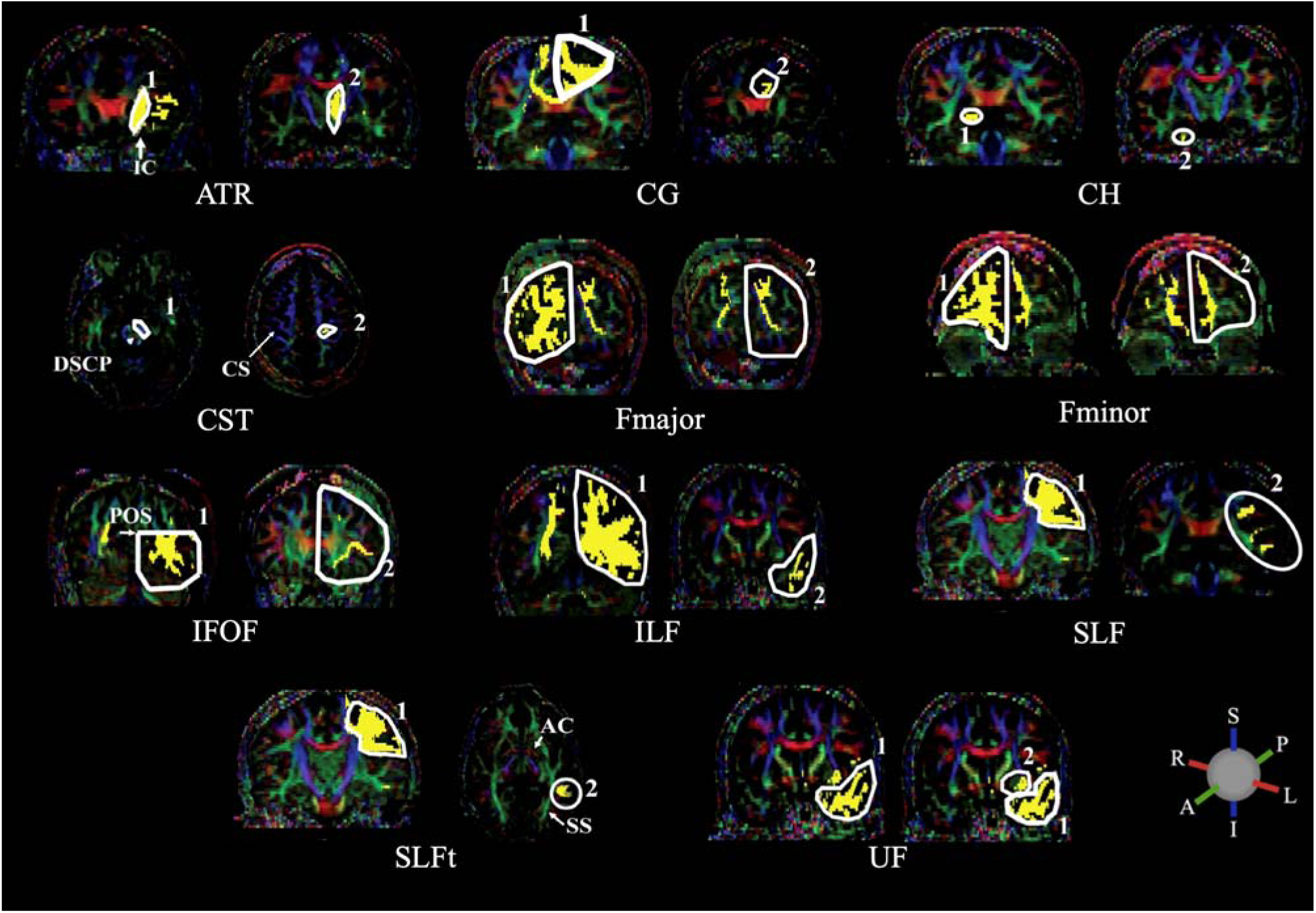
ROIs used to delineate the following major white matter tracts: anterior thalamic radiation (ATR); cingulate gyrus part of the cingulum (CG); hippocampal part of the cingulum (CH); cortico-spinal tract (CST); forceps major (Fmajor); forceps minor (Fminor); inferior fronto-occipital fasciculus (IFOF); inferior longitudinal fasciculus (ILF); superior longitudinal fasciculus (SLF); temporal part of the superior longitudinal fasciculus (SLFt); uncinate fasciculus (UF). Streamlines are included in a given tract if they pass through both 1 AND 2. The following abbreviations indicate anatomical landmarks used to draw the ROIs: internal capsule (IC); decussation of the superior cerebellar peduncle (DSCP); central sulcus (CS); parieto-occipital sulcus (POS); anterior commissure (AC); sagittal stratum (SS). ROIs are drawn in white with streamlines in yellow, overlaid on FA images with principal diffusion directions indicated by the colour ball; blue = superior-inferior (S-I), green = anterior-posterior (A-P) and red = right-left (L-R). Adapted from Hua et al., 2008, with permission from Elsevier.

To delineate the fornix, streamlines were included which pass through the body of the fornix and either of the posterior limbs which project to the hippocampus (Figure 3). These were isolated by first selecting an axial level at the lower edge of the splenium of the corpus callosum, as seen in the mid-sagittal plane (Figure 3, left); in this axial level, the first ROI was drawn around the body of the fornix. Viewing the streamlines which are delineated by the first ROI, additional bilateral ROIs were defined to include only those which project posteriorly from the fornix body (Figure 3, right).

**Figure 3.**
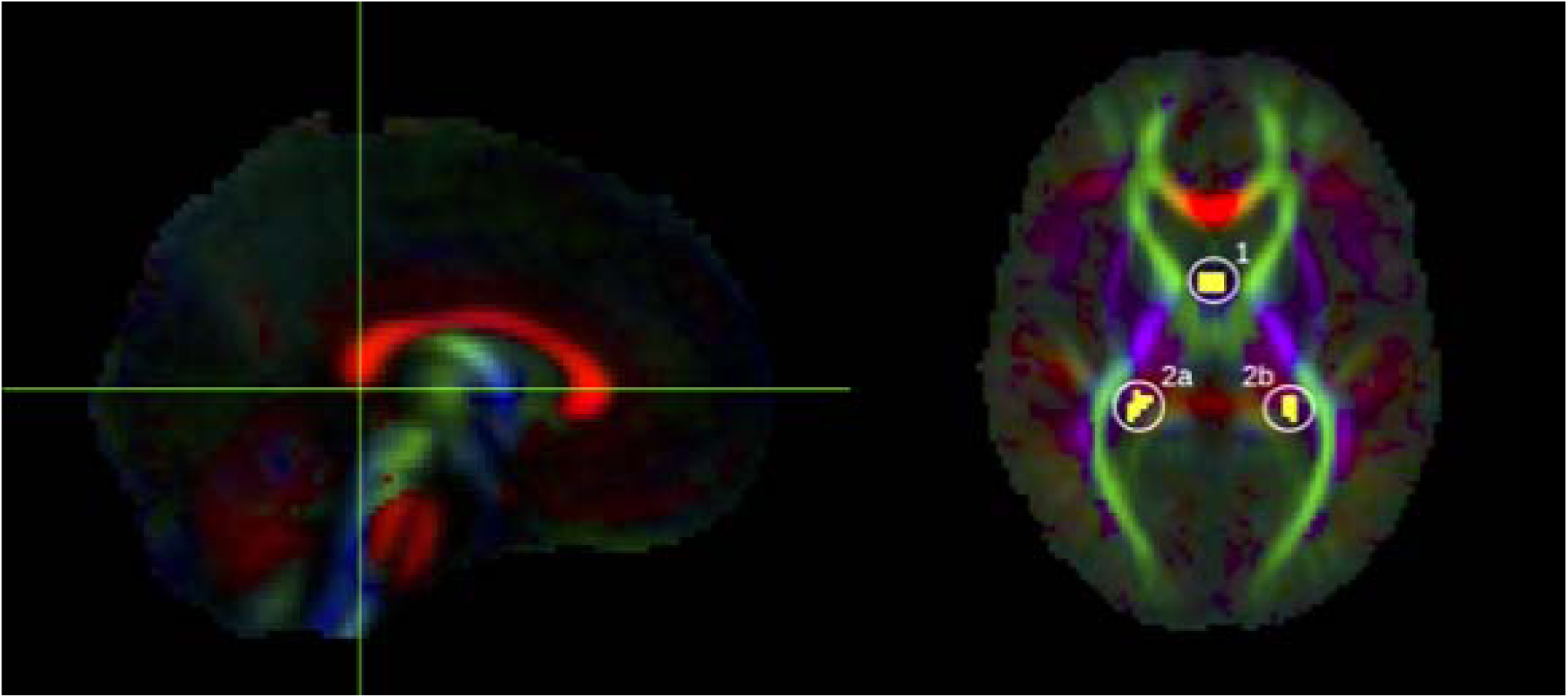
ROIs used to delineate the fornix, shown here on the group FA template. Yellow voxels contain streamlines which pass through the body of the fornix (1) AND bilateral posterior limbs of fornix (2a OR 2b).

For spatial normalisation, the average diffusion weighted image (aDWI), created for each subject by averaging all DWI images, was registered to the JHU aDWI template by 12-degree of freedom affine registration using FSL’s FLIRT (Jenkinson et al., 2002). The resulting transformation was then applied to the segmented streamlines. Any voxel containing one or more of these streamlines was then labelled, to create a binary mask for the tract for each individual. The average, across 28 subjects, of these binary masks was taken to give a probability map for each tract. The aDWI was then created for the group by averaging transformed aDWIs from all 28 subjects, and the group FA image was created from the group-average tensor map.

This atlas is available at Neurovault (https://neurovault.org/collections/LWTAAAST/).

### 2.5 Validation

The age-specific atlas was assessed by comparison with subject-specific tracts, delineated by applying the ROI-based tract-tracing method, described above, to each validation subject. These tracts were transformed to the atlas space by nonlinearly registering each subject’s FA image to the group FA template, constructed from the 28 atlas subjects, using FSL’s FNIRT (Andersson et al., 2007). We used three methods to assess accuracy of the atlas: i) volumetric overlap; ii) slice-wise correlation of FA measurements; and iii) correlation of whole-tract FA measurements. The same methods were also applied to the JHU atlas for comparison.

#### 2.5.1 Volumetric Overlap

To compare spatial similarity between normalised data we tested the volumetric overlap between the probabilistic atlas (age-specific or JHU) and each individually traced tract by measuring the Dice score (Dice, 1945) over a range of probability thresholds. The amount of volumetric overlap between the atlas data and the individually traced tract depends on both i) the quality of registration of the individual to the template, and ii) the agreement between the atlas data and the results from tractography in the individual. Thus, if the template is a closely matched target for registration, and if the underlying anatomy and diffusion process captured by the atlas is a good match to the validation subjects, we expect the Dice scores to be high.

#### 2.5.2 Slice-wise Correlation

We assessed the ability of the atlas to reproduce individually traced DWI metrics by calculating the mean FA in the tract in every slice along the major axis of each tract (coronal slices for tracts which project anterior/posterior; axial slices for tracts which project dorsal/ventral). In individually traced tracts, average FA was calculated by taking the mean FA in all masked voxels. In the probabilistic atlases (age-specific or JHU), the FA was weighted by the probability in each voxel using the following equation:

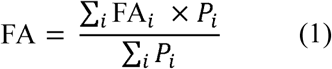

where FA_i_ is the FA in voxel i and P_i_ is the probability in voxel i. We then calculated the correlation between the probabilistic FA and individual FA (see Section 2.7).

#### 2.5.3 Whole-tract Correlation

Whole-tract average FA was calculated in each subject, using both probabilistic and individually traced tracts. Average FA was calculated in probabilistic tracts using equation (1) and in individually traced tracts by averaging FA in all masked voxels. We then calculated the correlation between the probabilistic FA and individual FA (see Section 2.7).

#### 2.5.4 Healthy Brain Network (HBN) Data

In order to alleviate bias associated with using same-site scans for validation, we used an additional validation dataset obtained from the Healthy Brain Network (HBN, http://fcon_1000.projects.nitrc.org/indi/cmi_healthy_brain_network/) (Alexander et al., 2017), a data-sharing biobank from the Child Mind Institute. Scans were obtained from 15 subjects, aged 6-8 years, from release 7.0 from the CitiGroup Cornell Brain Imaging Centre dataset. These multi-shell DWI data were acquired on a Siemens 3 tesla Prisma scanner using using an echo-planar pulse sequence with the following parameters: TE = 100.2 ms; TR = 3320 ms; 81 slices; 1.8 × 1.8 × 1.8 mm resolution; multi-band acceleration factor = 3; b = 1,000 s mm^-2^ and b = 2,000 s mm^-2^, each with 64 directions, and one b = 0 image. Preprocessing and quality control pipelines were applied as described above, followed by calculation of FODs using multi-shell multi-tissue constrained spherical deconvolution (Jeurissen et al., 2014) and tractography as described above. This allowed validation using subjects scanned in a different scanner, and with different scanning parameters. To further alleviate bias associated with using the same tractography algorithm for atlas construction and validation we also ran tractography in this cohort using a deterministic tensor-based algorithm (Basser et al., 2000), in addition to the FOD-based tractography algorithm described above. To give an overall indication of the accuracy of the atlas in these datasets, we applied the whole-tract correlation method described above. For completeness, in-depth results of the volumetric overlap and slice-wise correlation for the HBN data are given in the Supporting Information.

### 2.6 CoolMRI Study

As a demonstration, the age-specific atlas was used to investigate tract-level differences in white matter microstructure between the case and control children of the CoolMRI study. In each of the tracts delineated by the age-specific atlas, the average whole-tract FA was calculated for each individual using equation (1). We then tested for group differences in whole-tract FA. Bilateral tracts were tested separately. For comparison, we repeated the analysis using the JHU adult atlas. In the absence of ground truth, only a qualitative comparison of results obtained with the two atlases was performed.

### 2.7 Statistical Methods

To assess whether the age-specific atlas gave better volumetric agreement with individually traced tracts than the JHU adult atlas, we performed a two-tailed, paired t-test comparing the peak Dice scores.

In the slice-wise FA analysis and whole-tract FA analysis, we measured the correlation between atlas measurements and individual measurements using a repeated measures correlation coefficient (Bland and Altman, 1995), which uses an analysis of variance to calculate the correlation coefficient between residuals of the repeated measurements. This method was used in slice-wise FA analysis to calculate the correlation coefficient without variation due to different subjects, and in the whole-tract FA analysis to calculate the correlation coefficient without variation due to different tracts.

For each validation method, we compared the correlation coefficient given by the age-specific atlas with that given by the JHU adult atlas, by applying Fisher’s z-transform to each correlation coefficient and estimating the 95% confidence intervals of the difference between these z-transformed correlation coefficients. The confidence intervals were estimated with a percentile bootstrap method (Wilcox and Muska, 2002). In the slice-wise correlations, a moving block bootstrap method was used to account for the spatial dependence of repeated measurements in each subject (Politis and Romano, 1992).

In the CoolMRI demonstration, Mann-Whitney U tests were applied to test for differences in the median FA between cases and controls in each tract, with Bonferroni correction applied to correct for family-wise error. Significant results have corrected p < 0.05.

## 3 Results

### 3.1 Participant Demographics

The CoolMRI study recruited 51 children, without CP, treated with TH for NE at birth and 43 control children matched for age, sex and SES (Lee-Kelland et al., 2020). Of the recruited children, 7 cases and 4 controls did not want to undergo scanning. A further 4 cases had incomplete data due to movement during the scan. Quality control led to the rejection of a further 6 cases and 2 controls. One further case and one control were rejected due to incorrect image volume placement. This left 33 cases and 36 controls. These controls were split into 28 (15 male) for atlas construction and 8 (4 male) for validation. Data for each set of participants, as well as for HBN subjects, is shown in Table 1.

**Table 1.** Demographics of participants in the atlas dataset, same-site validation dataset, HBN validation dataset, and the CoolMRI dataset. Mean (range) is shown for age; median (range) is shown for SES and FSIQ in the CoolMRI cohort. Also shown are p-values of t-tests between atlas data and validation data for validation cohorts, and between cases and controls for the CoolMRI cohort. SES is defined as follows: A= upper middle class, B = middle class, C1 = lower middle class, C2 = skilled working class, D = working class, E = casual worker or unemployed.

|  | Atlas | Same-site Validation |  | HBN Validation |  | CoolMRI |  |  |
| --- | --- | --- | --- | --- | --- | --- | --- | --- |
|  |  |  | p |  | p | Cases | Controls | p |
| n = | 28 | 8 |  | 15 |  | 33 | 36 |  |
| Age | 7.0 (6.1-7.9) | 7.0 (6.1-7.8) | 0.9392 | 7.0 (6.0-7.9) | 0.8684 | 6.9 (6.0-7.9) | 7.0 (6.1-7.9) | 0.5595 |
| M/F | 15/13 | 4/4 | 0.8776 | 9/6 | 0.7002 | 18/15 | 19/17 | 0.8894 |
| SES |  |  |  |  |  | C1 (A-E) | B (A-D) | 0.1568 |
| FSIQ |  |  |  |  |  | 93 (62-115) | 108 (75-137) | <0.0001 |

### 3.2 Atlas

Figure 4 shows the probabilistic map for each tract, as well as the aDWI and FA images for the group of 28 children.

**Figure 4.**
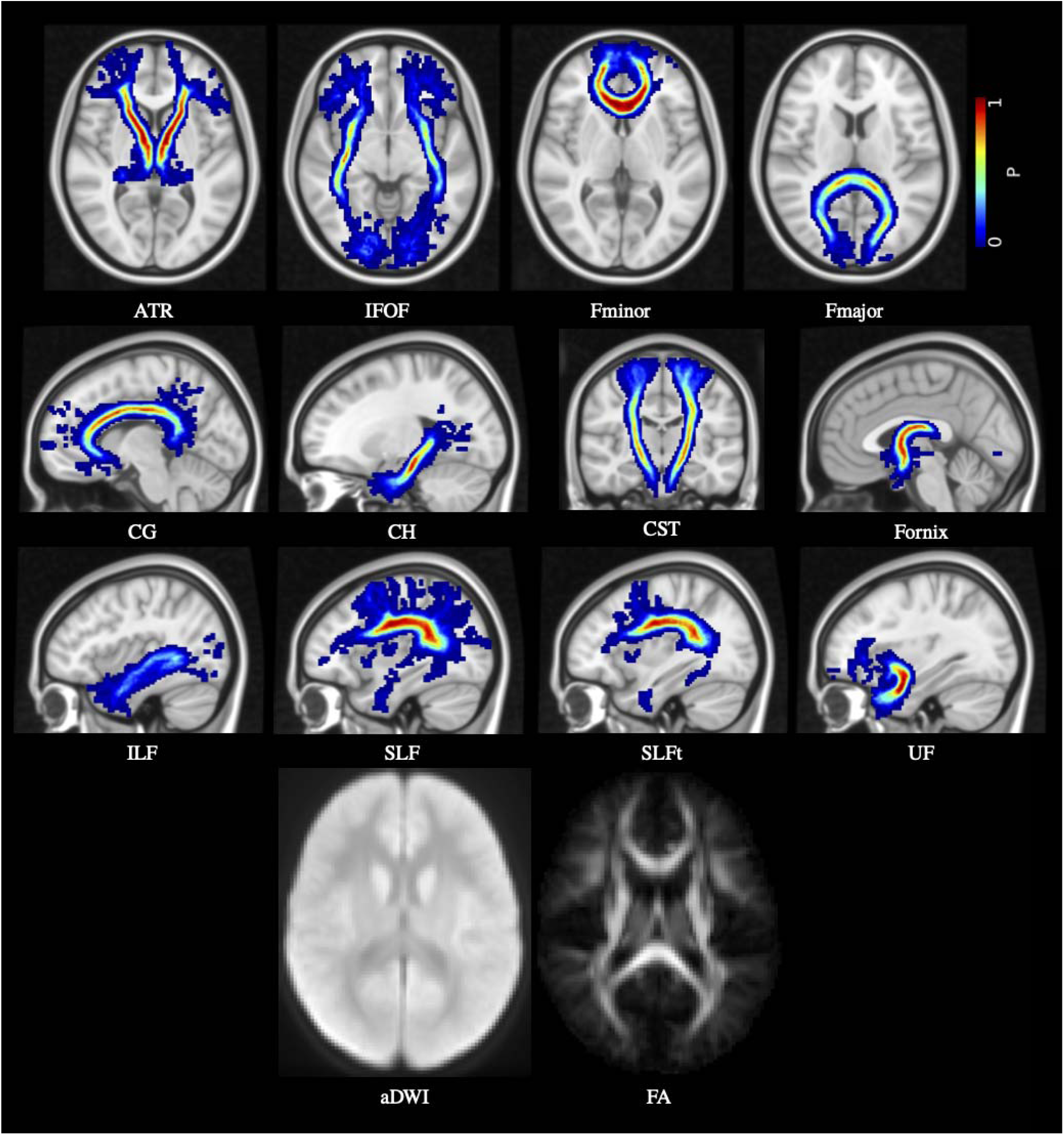
Age-specific probabilistic atlas for the 12 major white matter tracts: anterior thalamic radiation (ATR); inferior fronto-occipital fasciculus (IFOF); forceps minor (Fminor); forceps major (Fmajor); cingulate gyrus part of the cingulum (CG); hippocampal part of the cingulum (CH); cortico-spinal tract (CST); fornix; inferior longitudinal fasciculus (ILF); superior longitudinal fasciculus (SLF); temporal part of the superior longitudinal fasciculus (SLFt); and uncinate fasciculus (UF). Probabilities are indicated by the colour bar. Also shown are the aDWI and FA maps.

### 3.3 Validation

#### 3.3.1 Volumetric Overlap

The Dice score at a range of thresholds is plotted for each tract for the same-site validation data in Figure 5. The peak Dice scores for the age-specific atlas was significantly higher than for the JHU atlas in every tract (p < 0.05; see Table S1 for all p-values). The Dice scores for the HBN data are shown in Figures S1 and S2.

**Figure 5.**
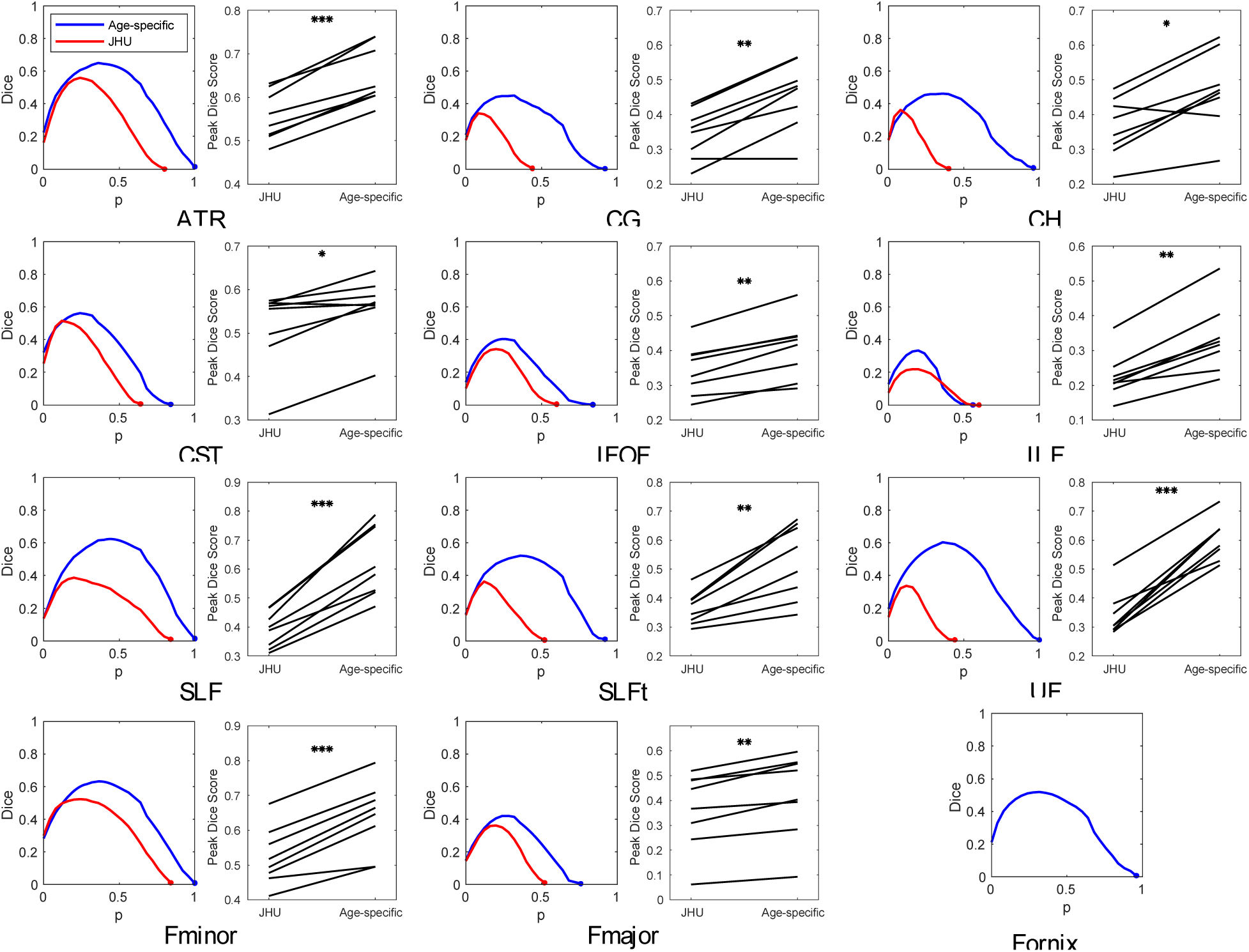
Same-site validation of tract overlap with “gold-standard” subject specific tract tracing. For each tract, the plot on the left shows the Dice score of volumetric overlap (y axis) against probability threshold (x axis) when using the age-specific atlas (blue) or the JHU adult atlas (red), with lines showing the mean score for the 8 validation subjects not included in the formation of the atlas, and shaded regions show the 95% confidence interval of the mean. Also shown for each tract is a paired plot of the peak Dice scores calculated with each atlas. P-values, given in Table S1, are indicated by: *p < 0.05; **p < 0.001; ***p < 0.0001. Note that the age-specific atlas outperformed the JHU (adult) atlas in all tracts. The tract representing the fornix is not available in the JHU atlas so only the new mask was tested.

#### 3.3.2 Slice-wise Correlation

The correlation between slice-wise FA measured by the age-specific atlas and that measured by subject-specific tract tracing is shown for the same-site validation data in Figure 6, with correlation coefficients measured using a repeated measures correlation (Bland and Altman, 1995). The correlations for the HBN data are shown in Figures S3 and S4. A correlation coefficient of one indicates perfect slice-wise agreement between the gold-standard (FA extracted from subject-specific tract tracing) and the FA estimated for each tract by registration to the either age-specific or JHU adult atlas. In the same-site data, most tracts showed strong correlation between FA measured by the age-specific atlas and that measured by subject-specific tract tracing, with all tracts having r > 0.8 apart from the CG (r = 0.625), SLF (r = 0.468) and SLFt (r = 0.546). The correlation coefficient for the age-specific atlas was higher than for the JHU adult atlas in all tracts, and this difference was significant in the ATR, CG, CST, Fminor and UF (indicated by the 95% confidence intervals of the difference between z-transformed correlation coefficients, see Table S2).

**Figure 6.**
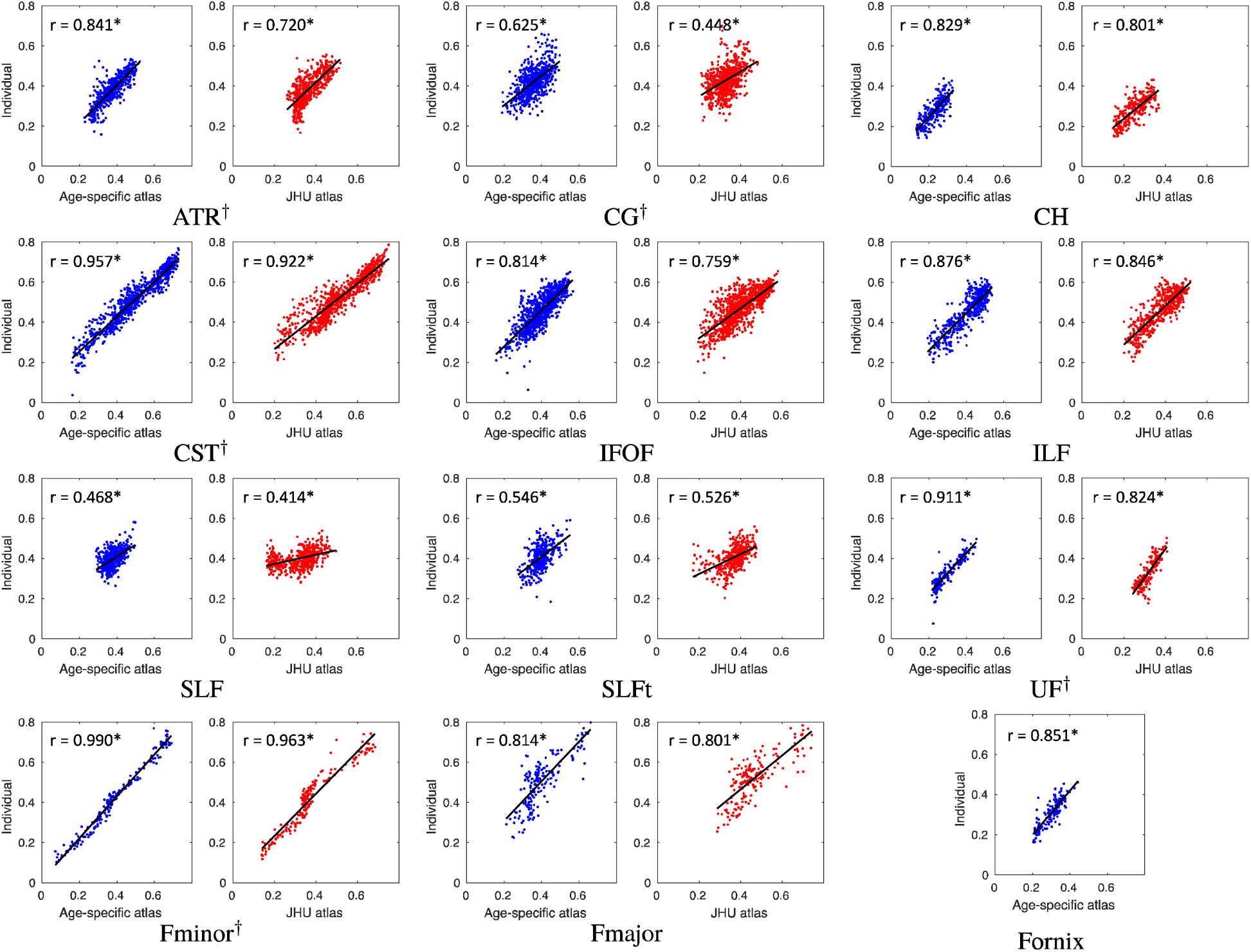
Same-site validation of slice FA values. Plots show slice FA measured from individually traced tracts (i.e. the “gold-standard”) plotted against corresponding values extracted from the age-specific and JHU atlases. Each plot shows a point for every slice in each of the 8 validation subjects and the regression. Correlation coefficients are shown on each plot, measured using a repeated measures correlation (Bland and Altman, 1995). All tracts exhibit higher correlation when measured with the age-specific atlas than with the JHU adult atlas. This difference is significant in the ATR, CG, CST, UF and Fminor, as indicated by † next to the tract abbreviation. Confidence intervals and regression parameters are given in Table S2. *p < 10^−20^.

#### 3.3.3 Whole-tract Correlation

Figure 7 shows the whole-tract FA measured by the atlas plotted against that given by subject-specific tract tracing for the same-site data, the HBN data with FOD-based tractography, and the HBN data with tensor-based tractography. The fornix is not included in these plots to allow valid comparison with the JHU atlas. Correlation coefficients, and confidence intervals of the difference between z-transformed correlation coefficients, are shown in Table 2. The age-specific atlas gave significantly stronger correlation of whole-tract FA measurements than the JHU adult atlas in all validation datasets.

**Table 2.** Validation of whole-tract FA correlations, corresponding to Figure 7. Columns show the parameters of the best-fit line y = mx + c and the correlation coefficient, r, between slice FA values from individual tracing and that from each atlas, measured using a repeated measures correlation (Bland and Altman, 1995). Also shown is the difference between the z- transform of the correlation coefficients for the age-specific atlas and the JHU atlas, and the 95% confidence intervals (CI) for this difference. Positive differences indicate a higher correlation with the age-specific atlas. These are shown for the same-site validation data, the HBN data with FOD-based tractography and the HBN data with tensor-based tractography. *p < 10^−10^.

|  | Age-specific Atlas |  |  | JHU Atlas |  |  | Difference between z-transformed correlation coefficients (95% CI) |
| --- | --- | --- | --- | --- | --- | --- | --- |
| Dataset | m | c | r | m | c | r |  |
| Same-site | 0.88 | 0.13 | 0.715* | 0.57 | 0.22 | 0.536* | +0.298 (+0.115, +0.300) |
| HBN (FOD) | 0.84 | 0.15 | 0.781* | 0.59 | 0.25 | 0.617* | +0.328 (+0.231, +0.412) |
| HBN (Tensor) | 0.51 | 0.27 | 0.697* | 0.39 | 0.32 | 0.595* | +0.176 (+0.087, +0.281) |

**Figure 7.**
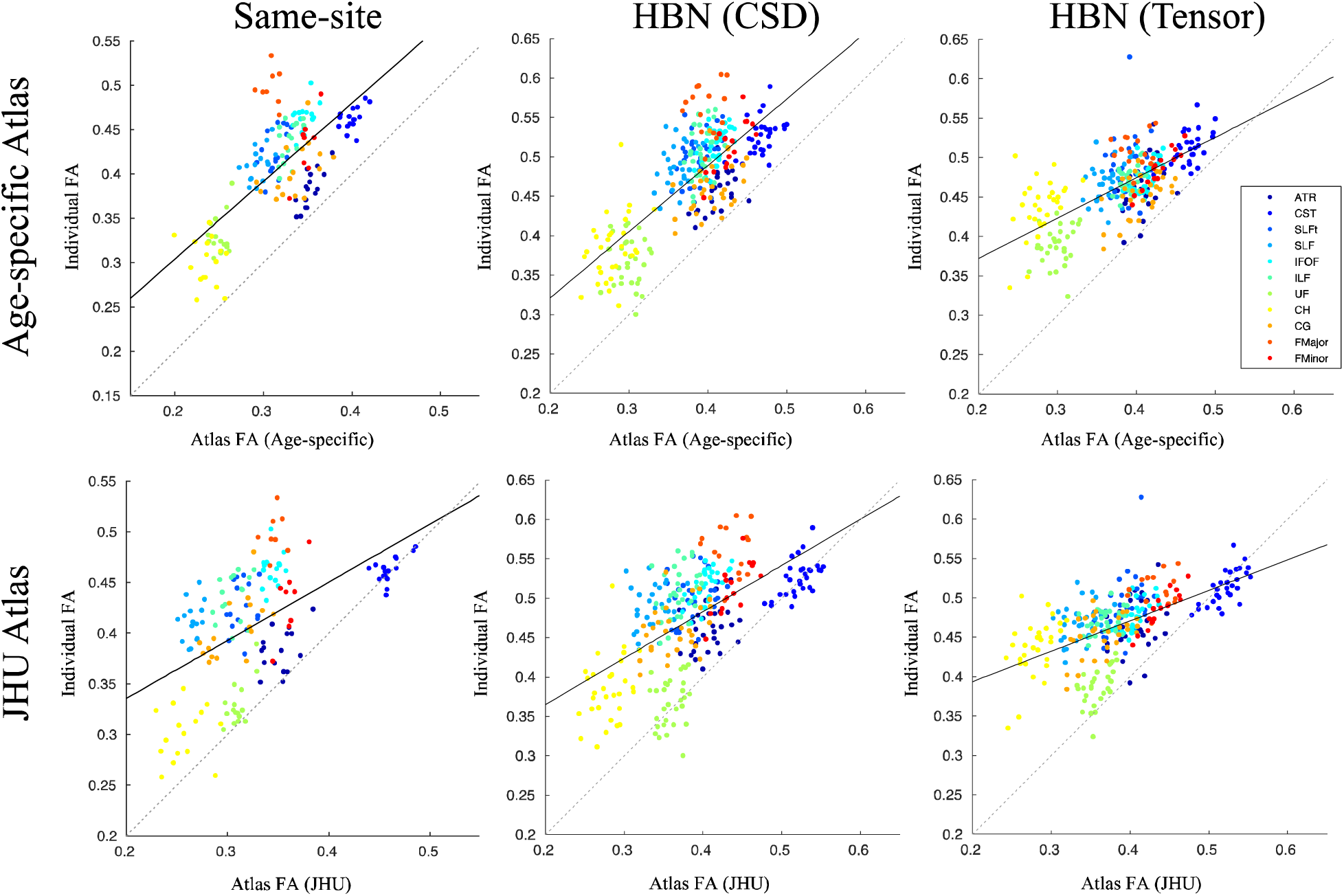
Comparison of mean FA extracted from whole tracts traced in individuals (“gold- standard”) and that estimated using each atlas. Whole-tract FA was measured by subject- specific tracing in the same-site validation data (left), the HBN data with FOD-based tractography (middle), and the HBN data with tensor-based tractography (right), then plotted against whole-tract FA measurements given by the age-specific atlas (top) or JHU adult atlas (bottom). The solid line shows the regression, and the dotted line represents exact equality between individual and the age-specific or JHU data. Correlation coefficients are given in Table 2.

### 3.4 CoolMRI Study

Numerous tracts in children treated with TH for NE had reduced FA compared to controls (see Table S5). After Bonferroni correction, only the left CG (p = 0.0056), left CH (p = 0.0081), left SLF (p = 0.0383), and fornix (p = 0.0121) had significantly reduced FA in cases compared to controls. The same analysis was run with the JHU atlas for comparison (see Table S6). Figure 8 shows box plots for both atlases for tracts in which at least one of the atlases revealed group differences in FA. Significant differences were found in the left SLF with the age-specific atlas but not the JHU adults atlas. Differences were found in the left CG and left CH with both atlases. Differences in the right CH were found with the JHU adult atlas but not with the age-specific atlas. Differences were found in the fornix with the age-specific atlas, but it is not available in the JHU atlas so could not be tested.

**Figure 8.**
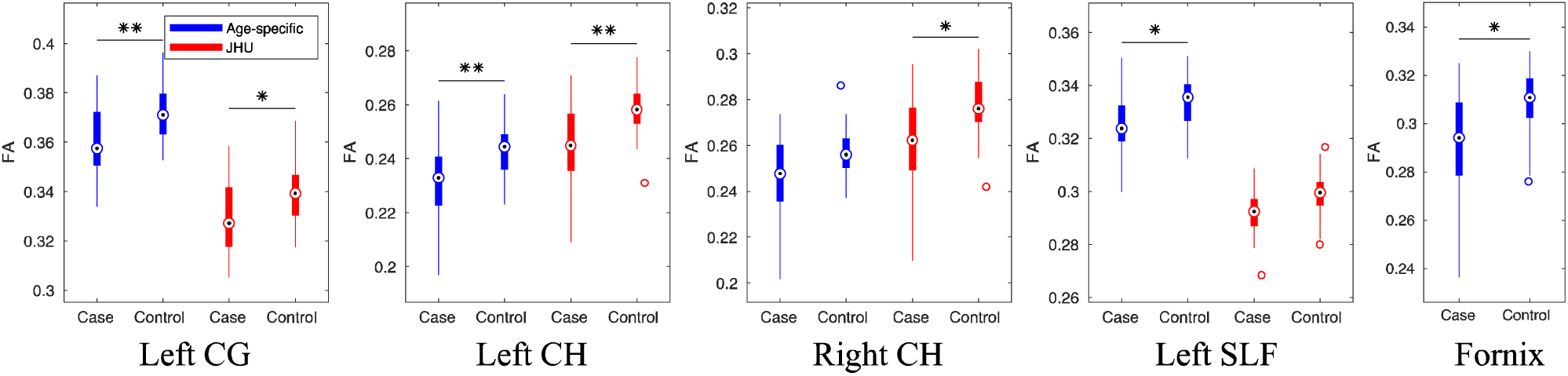
Box plots of significant differences in whole-tract average FA between children treated with TH for NE and healthy controls. Measurements obtained with both the age- specific atlas (blue) and the JHU adult atlas (red) are shown for tracts in which at least one of the atlases revealed significant differences between cases and controls; *p < 0.05, **p < 0.01, Bonferroni corrected. The fornix is not available in the JHU atlas so was only tested with the age-specific atlas. In each box, the point shows the median, the box shows the 25th to 75^th^ percentiles, and the lines extend to the maximum and minimum data points, excluding outliers which are displayed as circles.

## 4 Discussion

This study introduces an age-specific probabilistic white matter atlas constructed from children aged 6-8 years, providing a method of delineating white matter tracts without tractography. We have shown that this atlas accurately delineates tracts in children from a same-site cohort, and a cohort from a different site, imaged with different scanner and acquisition protocol. The strong correlation between FA sampled by the atlas and that measured in each individual (i.e. the “gold standard”), at both a whole-tract level and slice-wise level, shows that the atlas provides an accurate estimate for the underlying white matter microstructure. Additionally, the Dice scores between tracts in the atlas and those delineated by subject-specific tract tracing were higher with the age-specific atlas than with the JHU adult atlas, demonstrating improved anatomical accuracy of the age-specific atlas. In these validation methods, the age-specific atlas performed better than simply registering to an existing adult white matter tract atlas, as is routinely done in the literature. As a demonstration, we applied the age-specific atlas to the CoolMRI study, revealing significantly reduced FA in several major white matter tracts in children treated with TH for NE at birth compared to healthy controls.

The correlation of whole-tract FA measured by the atlas with that given by subject-specific tract tracing offers quantification of the performance of the atlas as a whole. In the same-site validation data, the HBN data traced with FOD-based tractography and the HBN data traced with tensor-based tractography, the age-specific atlas exhibited stronger correlation with the individual measurements than for the JHU atlas (Figure 7, Table 2). This shows that the age-specific atlas can accurately characterise the distribution of tract-level diffusion metrics in a study group, facilitating more sensitive group comparison and investigation of associations between these metrics and neuropsychological and behavioural measures.

Those tracts which exhibit low correlation between atlas and individual slice-wise FA measurements (namely the CG, SLF and SLFt) have very little spread in FA values, resulting in tightly grouped measurements with a low correlation coefficient (Figure 6). For these tracts, the Dice scores in Figure 5, as well as the tract-wise validation in Figure 7 demonstrate improved performance of the age-specific atlas at the level of whole tracts.

Long, thin tracts, such as the CST, IFOF and ILF, are particularly susceptible to partial volume effects when measuring volumetric overlap; a small radial translation will result in a large change to the Dice score. In these tracts, the high correlation in sampled FA values shows that the age-specific atlas gives accurate measurement of the tract microstructure.

Multi-site validation alleviates bias associated with using the same scanner for validation data and atlas construction, thus validation with the HBN data demonstrates that the age-specific atlas generalises to data from a different site, acquired with a different scanning protocol. In this dataset, the age-specific atlas gave better correlation of whole-tract FA measurements (Figure 7, Table 2). Additionally, the volumetric overlap in this dataset is significantly higher with the age-specific atlas than with the JHU adult atlas in all tracts apart from the CST and Fmajor, in which neither atlas performed significantly better than the other (Figure S1, Table S1). The age-specific atlas gave higher slice-wise correlations than the JHU adult atlas in all tracts; this difference was significant in the ATR, CST, Fminor, IFOF, ILF and UF (Figure S3, Table S3). There were no tests in which the JHU adult atlas performed significantly better in this dataset.

Further bias may be introduced by the use of the same tractography algorithm for both atlas generation and in estimating diffusion metrics for the validation data. Therefore, we also included a validation dataset in which subject-specific tract tracing was performed using a tensor-based tractography algorithm. Whereas the FOD-based tractography algorithm used to construct the age-specific atlas uses spherical deconvolution to find the peak FOD in the closest orientation to the propagating streamline, the tensor-based algorithm propagates the streamline along the principal eigenvector of the diffusion tensor at each step. This is comparable to the tensor-based tractography algorithm used in the construction of the JHU adult atlas, thus providing a conservative test case for validation. Despite this bias towards the JHU atlas, the age-specific atlas still gave stronger correlation of whole-tract FA measurements. In the tests of volumetric overlap (Figure S2, Table S1) and slice-wise correlation (Figure S4, Table S4) in this dataset, the age-specific atlas performed significantly better than the JHU adult atlas in at least one of these tests in six tracts (ATR, CH, ILF, UF, Fmajor, Fminor). In four tracts (CG, IFOF, SLF, SLFt) neither atlas performed significantly better in either test. In one tract (CST) the JHU atlas gives better volumetric overlap.

This introduces the question of how to provide the “gold-standard” of fibre tracking; the tensor-based algorithm was used in one of the HBN datasets in order to eliminate bias towards the age-specific atlas (by introducing bias towards the JHU adult atlas). However, this category of fibre tracking algorithm is widely acknowledged to give poor characterisation of diffusion in brain white matter due to its inability to resolve crossing fibres (Behrens et al., 2007; Tournier et al., 2012). Thus, the FOD-based algorithm used in the other validation datasets and in the construction of the atlas, which facilitates more comprehensive tracing due to its superior performance in regions of crossing fibres (Tournier et al., 2008), arguably gives a more accurate representation of the ground truth (i.e. the underlying white matter fibres). Therefore, when comparing the atlas to individually traced tracts in the validation data, the FOD-based algorithm likely gives a better indication of performance overall. Consequently, we believe the HBN data with tensor-based tract tracing provides a worst-case performance estimate, yet the age-specific atlas still out-performs the adult JHU atlas in many tests.

In future, as well as providing coverage of other age ranges, atlases could offer more extensive labelling of additional tracts and regions of white matter throughout development. A comprehensive database of traced tracts across a range of ages, potentially constructed by applying automated tractography-based white matter tract segmentation protocols (Lawes et al., 2008; Verhoeven et al., 2009; Wassermann et al., 2010; Zhang et al., 2018) to data from population studies such as the Human Connectome Project (Van Essen et al., 2013), Developing Human Connectome Project (Hughes et al., 2017), or Baby Connectome Project (Howell et al., 2019), would allow study-specific atlases to be built from subjects matched to a given study cohort.

Applying the age-specific atlas to the CoolMRI study to investigate group differences in tract-level FA revealed selective reduction in FA, that was significantly reduced in the left CG, left CH, left SLF and the fornix (Table S5). For comparison, we performed the same analysis with the JHU adult atlas (Table S6). Figure 8 demonstrates the differences in FA measurements from the different atlases. These differences result in some tracts exhibiting group differences in one atlas but not the other (right CH and left SLF). Due to the absence of ground truth, these results do not support the use of one atlas over another. However, these results demonstrate that the two atlases can give differing outcomes in a case-control study. Quantitative results from the validation methods above indicate that the age-specific atlas gives more accurate delineation of white matter tracts in this age group than the JHU adult atlas, suggesting the CoolMRI results obtained with the age-specific atlas are more reliable.

Previous studies of neonates treated with TH for NE have investigated the relationship between white matter diffusion properties, measured in the first weeks following birth, and neurodevelopmental outcome at 2 years of age. These studies found a significant reduction in FA in infants with adverse outcomes, compared to those with favourable outcomes, in widespread areas of white matter including, but not limited to the corpus callosum, anterior and posterior limbs of the internal capsule, external capsule, fornix, cingulum, and ILF (Lally et al., 2019; Tusor et al., 2012). Many of these regions were also shown to have reduced FA in the CoolMRI cases, indicating that the early structural alterations resulting from the brain injury cause lasting changes to white matter development. These results also provide evidence for an underlying white matter deficit which manifests as neuropsychological differences seen at school-age (Jary et al., 2019; Lee-Kelland et al., 2020; Tonks et al., 2019). Further investigation is required to link these structural impairments to specific components of the cognitive and motor assessments, and to develop therapeutic intervention strategies.

## 5 Conclusions

The age-specific atlas provided by this study has been shown to robustly delineate white matter tracts in children aged 6-8 years. Diffusion metrics sampled by the atlas correlate strongly with those measured by individual fibre tracking, allowing reliable investigation of white matter microstructure in cohorts. The closer agreement between FA measured in individually identified tracts and that estimated when registering to an age-specific atlas, suggests that such an approach would increase sensitivity to group differences, and is recommended for all studies performed in children.

## Data Availability Statement

This atlas is available at Neurovault (https://neurovault.org/collections/LWTAAAST/). The raw data that support the findings of this study are available from the corresponding author upon reasonable request.

## Ethics Approval Statement

Ethics approval was obtained from the North Bristol Research Ethics Committee and the Health Research Authority (REC ID: 15/SW/0148).

## Patient Consent Statement

Informed and written consent was obtained from the parents of participants before collecting data.

## Permissions

Permission was obtained from the relevant publisher for any reused content.

## Acknowledgements

We thank the children and their families for participating, Ngoc Jade Thai for the assistance with MR sequences, Aileen Wilson for her radiographical expertise, and James Tonks, Charlotte Whitfield and Emily Broadbridge for their assistance with neuropsychological assessment. We also thank Peter Green for statistical advice via the University of Bristol Statistics Clinic.

