## Supporting information for "An Age-Specific Atlas for Delineation of White Matter Pathways in Children Aged 6-8 Years"

**Figure S1:** HBN validation of tract overlap with “gold-standard” subject-specific tract tracing. For each tract, the plot on the left shows the Dice score of volumetric overlap (y axis) against probability threshold (x axis) when using the age-specific atlas (blue) or the JHU adult atlas (red), with lines showing the mean score for the 15 HBN subjects, and shaded regions show the 95% confidence interval of the mean. Also shown for each tract is a paired plot of the peak Dice scores calculated with each atlas. P-values, given in Table S1, are indicated by: *p < 0.05; **p < 0.001; ***p < 0.0001. Note that in all tracts apart from the CST and Fmajor, the age-specific atlas (derived from data acquired on a different scanner in different subjects with a different tractography algorithm) outperformed the JHU (adult) atlas. The tract representing the fornix is not available in the JHU atlas so only the new mask was tested.

**Figure S2:** HBN validation of tract overlap with “gold-standard” subject-specific tract tracing with a tensor-based tractography algorithm similar to that used to construct the JHU atlas. For each tract, the plot on the left shows the Dice score of volumetric overlap (y axis) against probability threshold (x axis) when using the age-specific atlas (blue) or the JHU adult atlas (red), with lines showing the mean score for the 15 HBN subjects, and shaded regions show the 95% confidence interval of the mean. Also shown for each tract is a paired plot of the peak Dice scores calculated with each atlas. P-values, given in Table S1, are indicated by: *p < 0.05; **p < 0.001; ***p < 0.0001. In the CH, ILF, UF and Fminor, the age-specific atlas performs better, whereas in the CST the JHU atlas performs better. The tract representing the fornix is not available in the JHU atlas so only the new mask was tested.

| Tract | Same-site | HBN (FOD) | HBN (Tensor) |
| --- | --- | --- | --- |
| ATR | <0.0001 | <0.0001 | n.s. |
| CG | 0.0006 | <0.0001 | n.s. |
| CH | 0.0031 | 0.0007 | 0.0008 |
| CST | 0.0093 | n.s. | 0.0003^†^ |
| Fmajor | 0.0007 | n.s. | n.s. |
| Fminor | <0.0001 | <0.0001 | <0.0001 |
| IFOF | 0.0001 | <0.0001 | n.s. |
| ILF | 0.0002 | <0.0001 | <0.0001 |
| SLF | 0.0001 | <0.0001 | n.s. |
| SLFt | 0.0010 | <0.0001 | n.s. |
| UF | <0.0001 | <0.0001 | 0.0160 |
| Fornix | - | - | - |

**Table S1:** p-values given by paired t-tests of the peak Dice scores measuring volumetric overlap between the atlas and the tracts delineated by subject-specific tracing. This is shown for the same-site data, the HBN data with FOD-based tractography and the HBN data with tensor-based tractography, corresponding to the results shown in Figure 5, Figure S1 and Figure S2 respectively. The Dice scores for the age-specific atlas are higher than those for the JHU atlas in all tests apart from the CST when traced with the tensor-based tractography algorithm in the HBN data, as indicated by †.

|  | Age-Specific Atlas | | | JHU Atlas | | | Difference between z-transformed correlation coefficients (95% CI) |
| --- | --- | --- | --- | --- | --- | --- | --- |
| Tract | m | c | r | m | c | r |  |
| ATR | 0.94 | 0.03 | 0.841* | 0.98 | 0.03 | 0.720* | +0.318 (+0.099, +0.548) |
| CG | 0.73 | 0.16 | 0.625* | 0.63 | 0.22 | 0.448* | +0.252 (+0.079, +0.428) |
| CH | 1.03 | 0.04 | 0.829* | 0.84 | 0.07 | 0.801* | +0.084 (-0.138, +0.289) |
| CST | 0.87 | 0.08 | 0.957* | 0.82 | 0.10 | 0.922* | +0.309 (+0.134, +0.461) |
| Fmajor | 0.98 | 0.11 | 0.814* | 0.84 | 0.13 | 0.801* | +0.040 (-0.376, +0.340) |
| Fminor | 1.04 | 0.01 | 0.990* | 1.05 | 0.02 | 0.963* | +0.669 (+0.363, +0.946) |
| IFOF | 0.90 | 0.10 | 0.814* | 0.75 | 0.17 | 0.759* | +0.144 (-0.001, +0.320) |
| ILF | 0.94 | 0.07 | 0.876* | 0.97 | 0.09 | 0.846* | +0.118 (-0.044, +0.282) |
| SLF | 0.58 | 0.18 | 0.468* | 0.23 | 0.33 | 0.414* | +0.068 (-0.077, +0.300) |
| SLFt | 0.71 | 0.12 | 0.546* | 0.50 | 0.22 | 0.526* | +0.029 (-0.223, +0.280) |
| UF | 1.02 | 0.02 | 0.911* | 1.42 | -0.12 | 0.824* | +0.367 (+0.189, +0.708) |
| Fornix | 1.11 | -0.02 | 0.851* | n/a | n/a | n/a | n/a |

**Table S2:** Validation of slice-wise FA measurements in the same-site validation data, corresponding to Figure 6 in the main text. Columns show the parameters of the best-fit line y = mx + c and the correlation coefficient, r, between slice FA values from individual tracing and that from each atlas, measured using a repeated measures correlation (Bland and Altman, 1995). Also shown is the difference between the z-transform of the correlation coefficients for the age-specific atlas and the JHU atlas, and the 95% confidence intervals (CI) for this difference. Positive differences indicate a higher correlation with the age-specific atlas. *p < 10^-20^.

**Figure S3:** HBN validation of slice FA values. Plots show slice FA measured from individually traced tracts (i.e. the “gold-standard”) plotted against corresponding values extracted from the age-specific and JHU atlases. Each plot shows a point for every slice in each of the 15 HBN subjects and the regression. All tracts exhibit higher correlation when measured with the age-specific atlas than with the JHU adult atlas. This difference is significant in the ATR, CST, IFOF, ILF, UF and Fminor, as indicated by † next to the tract abbreviation. Correlation coefficients are shown in Table S3.

|  | Age-Specific Atlas | | | JHU Atlas | | | Difference between z-transformed correlation coefficients (95% CI) |
| --- | --- | --- | --- | --- | --- | --- | --- |
| Tract | m | c | r | m | c | r |  |
| ATR | 1.03 | 0.02 | 0.743** | 0.80 | 0.15 | 0.642** | +0.195 (+0.031, +0.388) |
| CG | 0.78 | 0.13 | 0.602** | 0.72 | 0.21 | 0.563** | +0.059 (-0.045, +0.241) |
| CH | 0.62 | 0.17 | 0.440* | 0.50 | 0.21 | 0.331* | +0.129 (-0.091, +0.309) |
| CST | 0.84 | 0.12 | 0.925** | 0.78 | 0.15 | 0.895** | +0.175 (+0.067, +0.297) |
| Fmajor | 1.11 | 0.09 | 0.945** | 0.96 | 0.10 | 0.912** | +0.240 (-0.028, +0.417) |
| Fminor | 0.96 | 0.06 | 0.992** | 0.96 | 0.06 | 0.974** | +0.583 (+0.332, +0.777) |
| IFOF | 0.92 | 0.10 | 0.817** | 0.77 | 0.18 | 0.773** | +0.119 (+0.016, +0.222) |
| ILF | 0.90 | 0.12 | 0.741** | 0.70 | 0.24 | 0.622** | +0.223 (+0.080, +0.388) |
| SLF | 0.64 | 0.20 | 0.422** | 0.36 | 0.34 | 0.421** | +0.001 (-0.128, +0.171) |
| SLFt | 0.64 | 0.19 | 0.360** | 0.33 | 0.35 | 0.334* | +0.029 (-0.126, +0.199) |
| UF | 0.82 | 0.09 | 0.785** | 0.88 | 0.04 | 0.623** | +0.329 (+0.075, +0.479) |
| Fornix | 1.15 | -0.01 | 0.864** | n/a | n/a | n/a | n/a |

**Table S3:** Validation of slice-wise FA measurements in the HBN data, corresponding to Figure S3. Columns show the parameters of the best-fit line y = mx + c and the correlation coefficient, r, between slice FA values from individual tracing and that from each atlas, measured using a repeated measures correlation (Bland and Altman, 1995). Also shown is the difference between the z-transform of the correlation coefficients for the age-specific atlas and the JHU atlas, and the 95% confidence intervals for this difference. Positive differences indicate a higher correlation with the age-specific atlas. *p < 10^-6^, **p < 10^-20^.

**Figure S4:** HBN validation of slice FA values, using a tensor-based tractography algorithm similar to that used to construct the JHU atlas. Plots show slice FA measured from individually traced tracts (i.e. the “gold-standard”) plotted against corresponding values extracted from the age-specific and JHU atlases. Each plot shows a point for every slice in each of the 15 HBN subjects and the regression. The correlation is higher when measured with the age-specific atlas than with the JHU adult atlas for all tracts apart from the SLF and UF. This difference is significant in the ATR, ILF and Fmajor, as indicated by † next to the tract abbreviation. Correlation coefficients are shown in Table S4.

|  | Age-Specific Atlas | | | JHU Atlas | | | Difference between z-transformed correlation coefficients (95% CI) |
| --- | --- | --- | --- | --- | --- | --- | --- |
| Tract | m | c | r | m | c | r |  |
| ATR | 1.01 | 0.03 | 0.715** | 0.87 | 0.10 | 0.596** | +0.211 (+0.031, +0.390) |
| CG | 0.65 | 0.19 | 0.463** | 0.55 | 0.07 | 0.395** | +0.084 (-0.036, +0.254) |
| CH | 0.59 | 0.23 | 0.283* | 0.55 | 0.24 | 0.266* | +0.018 (-0.186, +0.242) |
| CST | 0.66 | 0.18 | 0.869** | 0.61 | 0.21 | 0.840** | +0.108 (-0.010, +0.233) |
| Fmajor | 1.04 | 0.07 | 0.967** | 0.92 | 0.07 | 0.942** | +0.290 (+0.038, +0.470) |
| Fminor | 0.69 | 0.15 | 0.929** | 0.67 | 0.15 | 0.896** | +0.197 (-0.242, +0.291) |
| IFOF | 0.84 | 0.09 | 0.879** | 0.77 | 0.14 | 0.861** | +0.072 (-0.046, +0.198) |
| ILF | 0.73 | 0.14 | 0.912** | 0.74 | 0.17 | 0.868** | +0.216 (+0.081, +0.338) |
| SLF | 0.49 | 0.25 | 0.280** | 0.45 | 0.29 | 0.375** | -0.106 (-0.238, +0.062) |
| SLFt | 0.50 | 0.25 | 0.164* | 0.48 | 0.27 | 0.156* | +0.008 (-0.131, +0.251) |
| UF | 0.74 | 0.14 | 0.657** | 0.84 | 0.07 | 0.710** | -0.100 (-0.270, +0.062) |
| Fornix | 0.58 | 0.22 | 0.727** | n/a | n/a | n/a | n/a |

**Table S4:** Validation of slice-wise FA measurements in the HBN data using tensor-based tractography for subject-specific tract tracing, corresponding to Figure S4. Columns show the parameters of the best-fit line y = mx + c and the correlation coefficient, r, between slice FA values from individual tracing and that from each atlas, measured using a repeated measures correlation (Bland and Altman, 1995). Also shown is the difference between the z-transform of the correlation coefficients for the age-specific atlas and the JHU atlas, and the 95% confidence intervals (CI) for this difference. Positive differences indicate a higher correlation with the age-specific atlas. *p < 0.005 **p < 10^-12^.

| Tract | Case | Control | Uncorrected p | Corrected p |
| --- | --- | --- | --- | --- |
| ATR-L | 0.352 | 0.361 | 0.0177 | - |
| ATR-R | 0.351 | 0.361 | 0.0145 | - |
| CG-L | 0.359 | 0.372 | 0.0003 | 0.0056 |
| CG-R | 0.327 | 0.337 | 0.0036 | - |
| CH-L | 0.232 | 0.244 | 0.0004 | 0.0081 |
| CH-R | 0.246 | 0.257 | 0.0087 | - |
| CST-L | 0.402 | 0.410 | 0.0140 | - |
| CST-R | 0.413 | 0.420 | 0.0371 | - |
| Fmajor | 0.345 | 0.352 | 0.0195 | - |
| Fminor | 0.366 | 0.374 | 0.0118 | - |
| Fornix | 0.292 | 0.309 | 0.0006 | 0.0121 |
| IFOF-L | 0.362 | 0.370 | 0.0054 | - |
| IFOF-R | 0.358 | 0.366 | 0.0052 | - |
| ILF-L | 0.348 | 0.355 | 0.0340 | - |
| ILF-R | 0.350 | 0.356 | 0.0236 | - |
| SLF-L | 0.326 | 0.334 | 0.0018 | 0.0383 |
| SLF-R | 0.313 | 0.318 | - | - |
| SLFt-L | 0.344 | 0.352 | 0.0054 | - |
| SLFt-R | 0.331 | 0.335 | - | - |
| UF-L | 0.273 | 0.277 | - | - |
| UF-R | 0.269 | 0.271 | - | - |

**Table S5:** Comparison of mean FA in each tract in cases and controls using the age-specific atlas. Columns show mean of each group, the uncorrected p-values calculated by Mann-Whitney U test, and corrected p-values with Bonferroni correction applied.

| Tract | Case | Control | Uncorrected p | Corrected p |
| --- | --- | --- | --- | --- |
| ATR-L | 0.363 | 0.371 | 0.0177 | - |
| ATR-R | 0.341 | 0.350 | 0.0084 | - |
| CG-L | 0.330 | 0.340 | 0.0020 | 0.0416 |
| CG-R | 0.296 | 0.301 | - | - |
| CH-L | 0.245 | 0.258 | 0.0003 | 0.0059 |
| CH-R | 0.262 | 0.277 | 0.0009 | 0.0180 |
| CST-L | 0.458 | 0.466 | 0.0371 | - |
| CST-R | 0.461 | 0.467 | - | - |
| Fmajor | 0.381 | 0.385 | - | - |
| Fminor | 0.372 | 0.380 | 0.0166 | - |
| Fornix | n/a | n/a | n/a | n/a |
| IFOF-L | 0.353 | 0.360 | 0.0075 | - |
| IFOF-R | 0.358 | 0.365 | 0.0084 | - |
| ILF-L | 0.317 | 0.324 | 0.0123 | - |
| ILF-R | 0.332 | 0.338 | 0.0107 | - |
| SLF-L | 0.292 | 0.298 | 0.0061 | - |
| SLF-R | 0.301 | 0.304 | - | - |
| SLFt-L | 0.331 | 0.339 | 0.0052 | - |
| SLFt-R | 0.348 | 0.350 | - | - |
| UF-L | 0.319 | 0.324 | - | - |
| UF-R | 0.318 | 0.324 | - | - |

**Table S6:** Comparison of mean FA in each tract in cases and controls using the JHU atlas. Columns show mean of each group, the uncorrected p-values calculated by Mann-Whitney U test, and corrected p-values with Bonferroni correction applied. Note that Bonferroni correction is applied across 21 tracts to allow comparison of corrected p-values with those from the age-specific atlas.
